# Predictive steering: Integration of artificial motor signals in self-motion estimation

**DOI:** 10.1101/2022.06.03.494696

**Authors:** Milou J.L. van Helvert, Luc P.J. Selen, Robert J. van Beers, W. Pieter Medendorp

## Abstract

The brain’s computations for active and passive self-motion estimation can be unified with a single model that optimally combines vestibular and visual signals with sensory predictions based on motor efference copies. It is unknown whether this theoretical framework also applies to the integration of artificial motor signals, like the motor signals that occur when driving a car. Here, we examined if training humans to control a self-motion platform would lead to the construction of an accurate internal model of the mapping between the steering movement and the vestibular reafference. Participants (n = 15) were seated on a linear motion platform and actively controlled the platform’s velocity using a steering wheel to translate their body to a memorized visual target location (Motion condition). We compared their steering behavior to that of participants (n = 15) who remained stationary and instead aligned a non-visible line with the target (Stationary condition). To probe learning, the gain between the steering wheel angle and the platform velocity or line velocity changed abruptly twice during the experiment. These gain changes were virtually undetectable in the displacement error in the Motion condition, whereas clear deviations were observed in the Stationary condition. These results show that participants in the Motion condition made within-trial changes to their steering behavior immediately after the gain changes. This suggests that they continuously compared the vestibular reafference to internal predictions, and thus employed and updated an internal forward model of the mapping between the steering movement and the vestibular reafference.

**New & Noteworthy:** Perception of self-motion is known to depend on the integration of sensory signals and, when the motion is self-generated, the predicted sensory reafference based on motor efference copies. Here we show, using a closed-loop steering experiment with a direct coupling between the steering movement and the vestibular self-motion feedback, that humans are also able to integrate artificial motor signals, like the motor signals that occur when driving a car.

## Introduction

Self-motion estimation depends on the integration of sensory and motor information. During passively generated motion (e.g., a passenger in a moving car), perception of self-motion comes primarily from the visual system, which provides optic flow cues (1), and the vestibular system (2, 3). Because sensory signals may be ambiguous (e.g., the otoliths cannot distinguish between translational motion and gravitational acceleration), the brain is thought to use an internal sensory integration model that combines sensory information from different modalities to form a final self-motion percept (4–6).

When the motion is generated actively, the brain can also integrate information related to the motor command to estimate self-motion (see Ref. 7 for a review). In fact, self-motion is judged better when it is actively generated than passively imposed (8–11). Also, patients with vestibular deficits perceive self-motion significantly better when self-generated (12–15).

While these findings could be interpreted as evidence that vestibular signals (and sensory signals more generally) are functionally less important in actively moving subjects, recent modeling work has provided a unified theory for how active and passive motion can be estimated (16, 17), with a fundamental role for both sensory signals and the efference copy. According to this theory, a multisensory self-motion estimate is computed using sensory prediction errors, i.e., the difference between actual and predicted sensory signals. During active motion, motor commands can be used to anticipate the corresponding sensory reafference, such that the sensory prediction error is minimal. In contrast, sensory activity cannot be anticipated during passive motion, resulting in non-zero sensory prediction errors, which then drive the self-motion estimate.

Under both active and passive motion, vestibular signals (as well as other sensory signals like vision) are continuously monitored to update the internal prediction. Thus, without intact sensory organs, the sensory prediction errors cannot be corrected, and the self-motion estimate may no longer be accurate during either active or passive motion. Because sensory information and motor commands, as well as the neural processing itself, are endowed with intrinsic random noise, Laurens and Angelaki (16) modelled the computations using a Kalman filter to determine the optimal (Bayesian) estimate of self-motion. Given uncertainty in the moment-to-moment sensory information, such a Bayesian computation also relies on a priori expectations about incoming sensory signals (6, 18–20).

While this framework suggests that not only sensory signals but also efference copies of motor commands are critical in self-motion perception, it is agnostic as to the nature of the motor signal. This opens up the possibility that also artificial motor signals can be used for self-motion perception, as long as they are associated with an accurate internal model for predicting the sensory reafference. Such artificial motor signals are for example generated when driving a car; the steering is cognitively mediated and of efferent nature. The use of such artificial motor signals for self-motion perception is the topic of the present study.

Data on this issue are sparse and contradictory. For example, Roy and Cullen (21) taught monkeys to drive themselves using a steering wheel that controlled the speed of the turntable on which they were seated. They compared neural activity between an active steering condition and voluntary head rotation conditions. While neuronal activity was suppressed at early sensory levels during active head rotations, reflecting a near-zero prediction error, this was not observed during self-generated driving, during which neurons responded as if the motion was externally applied. These findings suggest that an artificial motor signal, here a cognitive steering signal, is not used to predict the sensory afference at early sensory levels. In contrast, other work (22, 23) has reported that neurons in the dorsal stream (medial superior temporal area) show altered responses to visual self-motion when monkeys steer to move in a certain direction compared to when they passively view the same optic flow pattern (but see also Ref. 24), as if the brain not only relied on sensory self-motion information but also made an internal model prediction based on steering-related signals.

In the present study we address this issue in humans, testing whether and how steering-related signals are used in self-motion perception. Recent virtual reality experiments examined how humans (and monkeys) virtually navigated to a memorized location by integrating optic flow generated by their own joystick movements (25–27). Biases in their steering depended on optic flow density, as a marker of the reliability of sensory evidence, and the control gain of the joystick, as a measure of the internal model prediction of the optic flow, suggesting that the brain combined both signals in the percept of non-vestibular self-motion. However, the authors mainly focused on the processing of visual information, and the role of the vestibular sense was only studied under continuously changing control dynamics of the joystick (27). How the brain formed an internal model to predict the vestibular self-motion signal thus remains unknown.

We created a motor signal of cognitive nature (an artificial efference copy) and test how it is used in combination with vestibular-derived self-motion signals. This outflow signal was generated by training subjects to drive their own body, by handling a steering wheel that controlled the lateral motion velocity of a vestibular platform, to a memorized visual target (Motion condition). We examined whether participants anticipate the sensory consequences of the steering induced motion and integrate these with the vestibular feedback signals. We focused on the dynamics by which participants learn and adjust their internal predictions and estimate of the platform motion by abruptly changing the gain between the steering wheel angle and the velocity of the platform twice during the experiment. If participants construct an internal model of the mapping between the steering movement and the vestibular reafference, we expect rapid, within-trial, changes to their steering behavior after these gain changes in order to align their body with the memorized target. We compared their behavior to that of participants who did not have vestibular feedback about their motion, and could thus only employ a feedforward strategy, as they handled the steering wheel to control a line cursor (Stationary condition).

## Methods

### Participants

Thirty participants were randomly assigned to one of two experimental conditions. The Motion group included 15 participants (five men and ten women) ranging in age from 18 to 35 years, and the Stationary group included 15 participants (six men and nine women) ranging in age from 18 to 29 years. All participants were naïve to the purpose of the experiment and reported to have normal or corrected-to-normal vision and no history of motion sickness. The ethics committee of the Faculty of Social Sciences of Radboud University Nijmegen, the Netherlands, approved the study and all participants gave written informed consent prior to the start of the study. Participants were reimbursed for their time with course credit or €12,50. The experimental session took around 75 minutes per participant.

### Setup

The experiment took place in a dark room. Participants were seated on a custom-built linear motion platform, also called the sled, with their interaural axis aligned with the motion axis of the sled (Fig. 1A). The track of the sled was approximately 95 cm long. The sled was powered by a linear motor (TB15N; Tecnotion, Almelo, The Netherlands) and controlled by a servo drive (Kollmorgen S700; Danaher, Washington, DC, United States). Participants were restrained by a five-point seat belt and could stop the motion of the sled at any time by pressing one of the emergency buttons on either side of the sled chair. A steering wheel (G27 Racing Wheel; Logitech, Lausanne, Switzerland) with a range of rotation from −450 to +450 deg and a resolution of 0.0549 deg was mounted in front of the participants at chest level. The steering wheel was placed at a comfortable handling distance from the body for each individual participant. The angle of the steering wheel encoded the linear velocity of the sled (Motion condition) or a vertical line cursor (Stationary condition). Visual stimuli were presented on a 55 inch OLED screen (55EA8809-ZC; LG, Seoul, South Korea) with a resolution of 1920 x 1080 pixels and a refresh rate of 60 Hz, positioned centrally in front of the sled track at a viewing distance of approximately 170 cm. Participants wore headphones during the entire experiment to mask the noise of the moving sled with white noise sounds. The experiment was controlled using custom-written software in Python (version 3.6.9; Python Software Foundation, https://www.python.org/).

**Figure 1.**
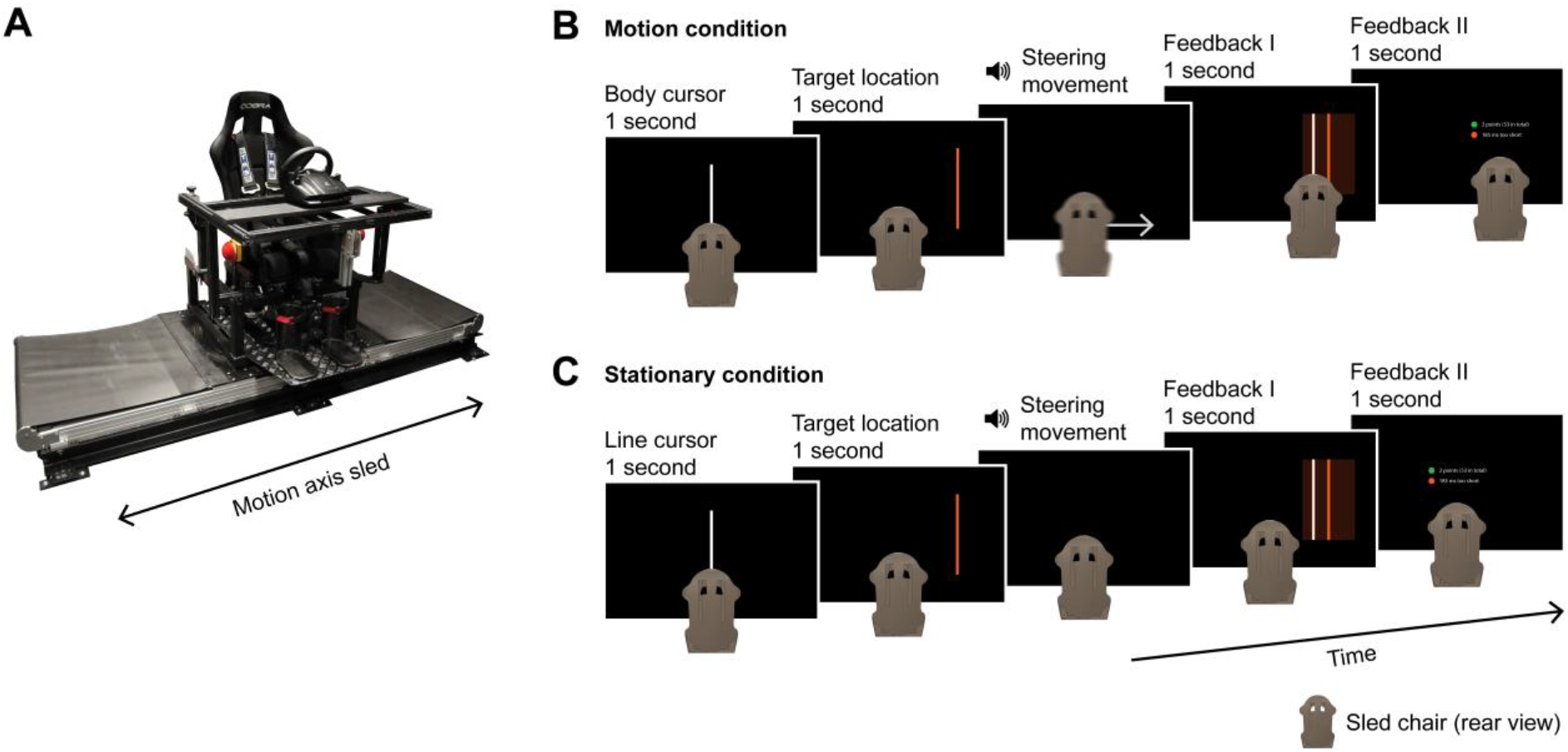
Experimental setup and paradigm. A) Experimental setup. Participants were seated with their interaural axis aligned with the motion axis of the sled and turned a steering wheel. B) Motion condition paradigm. Participants were first shown the body cursor (white line), followed by the target (orange line). After the disappearance of the target, a beep instructed participants to turn the steering wheel to translate their body in alignment with the memorized target location. After the motion, visual feedback about the distance from the reappearing body cursor to the target location (Feedback I) and the movement duration (Feedback II) was provided. C) Stationary condition paradigm. Participants were first shown the line cursor (white line), followed by the target (orange line). After the disappearance of the target, a beep instructed participants to turn the steering wheel to translate the memorized line cursor in alignment with the memorized target location. Participants remained stationary and did not receive any visual feedback during the steering movement. After the movement, visual feedback about the distance from the reappearing line cursor to the target location (Feedback I) and the movement duration (Feedback II) was provided.

### Paradigm

#### Motion condition

In the Motion condition, participants turned the steering wheel to laterally translate their body to align with a memorized visual target. The angle of the steering wheel encoded the linear velocity of the sled. The experimental session started with a two-minute familiarization with visual feedback to become acquainted with the initial gain between the angle of the steering wheel and the velocity of the sled (1.4 cm/s per deg, see below). After the familiarization, the main experiment started.

Figure 1B shows the sequence of events during an experimental trial. At the start of the trial, the position of the body midline was presented on the screen as a vertical white line with a length of 25.4 cm for 1 s. We will refer to this line as the body cursor. Next, the target, represented by a vertical orange line with the same length, was presented for 1 s. The target distance, defined as the distance from the body cursor to the target location, was 20, 30 or 40 cm. The target could appear to the left or to the right of the body midline. After disappearance of the target, a beep was played via the headphones to inform the participant to start the steering movement to align their body midline with the memorized target location.

The motion started when the participant turned the steering wheel 0.0549 deg (one “click”) away from the steering wheel angle at trial start. Participants received no visual information during the motion. As described above, the initial gain between the angle of the steering wheel and the velocity of the sled was 1.4 cm/s per deg. To probe learning, the gain changed abruptly twice during the experiment (trial 1-90: 1.4 cm/s per deg; trial 91-162: 0.8 cm/s per deg; trial 163-234: 1.4 cm/s per deg). Participants were not informed about the initial gain or the gain changes, and were instructed to make a smooth steering movement. The latency between the rotation of the steering wheel and the translation of the sled was typically lower than 10 ms. The maximum absolute velocity of the sled was set to 100 cm/s. If the steering wheel angle encoded a higher sled velocity, it was capped at this maximum velocity (< 1 trial per participant). During the motion, white noise was played through the headphones to mask any auditory cues. When the absolute velocity encoded by the steering wheel angle fell below 2 cm/s the sled stopped, and the white noise sound ended.

After the motion, participants received feedback about the accuracy of their displacement and the duration of the steering movement. First, both the body cursor and the target were presented on the screen for 1 s. This informed participants about how far they ended from the target location, and whether they undershot or overshot the target location with their self-generated motion. To incentivize participants to adequately perform the task they also received a score. Two points were awarded if the undershoot or overshoot was smaller than 0.15 times the target distance, represented on the screen by a translucent orange rectangular area centered on the target stimulus. One point or zero points were awarded if the undershoot or overshoot was between 0.15 and 0.30 times or larger than 0.30 times the target distance, respectively. Subsequently, a line of text reiterating the score and the total score so far and a line of text with the movement duration were presented on the screen for 1 s. Participants were encouraged to finish their steering movement within 900 to 1300 ms from movement start to ensure suprathreshold vestibular stimulation while remaining below the maximal sled velocity. The line of text read: “Timing perfect” if the movement ended after 900 to 1300 ms, and *“n* ms too short/long” if the movement took shorter or longer. The lines of text were preceded by colored circles, with the color quickly informing the participants about their performance (displacement accuracy: green, orange and red for two, one and zero points, respectively; movement duration: green, orange and red for a perfect timing, 300 ms too short or long and more than 300 ms too short or long, respectively).

Trials were presented in blocks of six trials with the target presented at different locations: 20, 30 and 40 cm to the left and right of the body cursor at trial start. Target distances within a trial block were presented in a semi-random order, with leftward and rightward displacements alternating, and each distance presented once in either direction. The sled started a trial at the location where the previous trial ended. However, if the position of the sled at the end of a trial was restricting its motion on the next trial (because of the limited sled track length of ~95 cm) to less than 1.5 times the target distance, the sled was first passively moved to a position 30 cm away from the middle of the sled track in the direction opposite that of the upcoming displacement, leaving ~80 cm for the motion. The main experiment started with 18 practice trials, during which the experimenter was present for task instructions. The practice trials were followed by the 234 experimental trials, of which the first always tested a rightward displacement. The experimental trials were separated by short breaks (< 2 minutes) after every 36 trials, during which the lights in the experimental room were turned on to prevent dark adaptation.

#### Stationary condition

In the Stationary condition, participants turned the steering wheel to laterally translate a non-visible line cursor in alignment with a memorized visual target, while the sled (and thus the body) remained stationary. The experimental session started with a two-minute familiarization with visual feedback to become acquainted with the initial gain between the angle of the steering wheel and the velocity of the line cursor (1.4 cm/s per deg, see below). After the familiarization, the main experiment started.

During the main experiment, targets were presented as in the Motion condition (Fig. 1C). However, instead of the body cursor, participants controlled a non-visible line cursor that moved independently of the stationary body. At the start of the trial, the line cursor was presented on the screen in front of the participant, aligned with the body midline, as a vertical white line with a length of 25.4 cm for 1 s. After the subsequent presentation of the target, a beep was played via the headphones to inform the participant to start the steering movement to align the line cursor with the memorized target. Note that neither the line cursor nor the target was visible during the steering movement. White noise was played through the headphones during the steering movement to keep conditions similar.

The gain between the steering wheel angle from trial start and the velocity of the line cursor changed over trials in the same way as in the Motion condition (trial 1-90: 1.4 cm/s per deg; trial 91-162: 0.8 cm/s per deg; trial 163-234: 1.4 cm/s per deg). During the steering movement, the position of the line cursor was updated in the background by adding up the products of the encoded velocities and the time between steering wheel samples. Contrary to the Motion condition, no maximum absolute velocity was set. When the absolute velocity encoded by the steering wheel angle fell below 2 cm/s the white noise sound ended and participants received feedback and a score as in the Motion condition (the updated position of the line cursor, in contrast to the body cursor, was shown along with the target).

Trials were presented in blocks of six trials as in the Motion condition. The main experiment started with 18 practice trials, during which the experimenter was present for task instructions. The practice trials were followed by the 234 experimental trials, of which the first always tested a rightward displacement. The experimental trials were separated by short breaks (< 2 minutes) after every 36 trials, during which the lights in the experimental room were turned on to prevent dark adaptation.

### Data analysis

Data were processed offline in MATLAB (version R2017a; The MathWorks, Inc., Natick, Massachusetts, United States). Trials during which participants displaced the sled (Motion condition) or the line cursor (Stationary condition) in the direction opposite of the target or during which participants rotated the steering wheel less than 7.5 deg from the angle at trial start were excluded from the analysis. Additionally, for the Motion condition, trials during which the absolute velocity encoded by the steering wheel angle reached the set maximum of 100 cm/s or during which the sled reached one of the ends of the sled track were excluded. On average, one trial was excluded per participant (mean ± SD; Motion condition: 1.40 ± 1.45 trials per participant; Stationary condition: 0.67 ± 0.82 trials per participant).

For all included trials movement onset was defined as the first time point the steering wheel rotated more than 2.5 deg away from the angle at trial start. Movement end was defined as the first time point after movement onset the steering wheel angle fell below 2.5 deg from the angle at trial start. Participants failed to bring the steering wheel angle back within this range (i.e., stopped steering prematurely) on average on five trials per participant (Motion condition: 4.60 ± 4.17 trials; Stationary condition: 5.20 ± 6.70 trials). For these trials, movement end was defined as the time point the steering wheel angle remained constant for at least 100 ms or reached a local minimum while encoding a low velocity (i.e., rotated less than 7.5 deg away from the angle at trial start). Movement duration was defined as the time between movement onset and movement end. Displacement error was defined as the distance between the body cursor (Motion condition) or the line cursor (Stationary condition) at movement end and the target. Negative errors represent undershoots; positive errors represent overshoots. Relative displacement errors were computed as the ratio of the displacement error and the target distance.

#### Normalized steering behavior

To be able to depict changes in steering behavior in response to the two gain changes across participants, we first normalized the time traces of the steering wheel angle. For each participant, we first calculated the mean movement duration and mean maximum absolute steering wheel angle of the baseline trials (trials 73-90, the last three trial blocks before the first gain change), grouped based on target distance and direction. We subsequently normalized the movement duration and steering wheel angle samples on each trial by dividing them by the mean movement duration and the mean maximum absolute steering wheel angle, respectively, of the three baseline trials with a corresponding target distance and direction. Normalized steering wheel angles were resampled to 1000 samples per trial using linear interpolation and were averaged across participants. We then created a corresponding linearly spaced time vector of 1000 samples for each trial running from zero, representing movement onset, to the mean normalized movement duration across participants for plotting purposes.

#### Scale factors and skewness

To quantify changes in steering kinematics in response to the two gain changes, we scaled both the raw time and raw steering wheel angle samples on each trial relative to the baseline trials with a corresponding target distance and direction. This linear transformation from baseline trial *b* to trial of interest *i* can be described by:

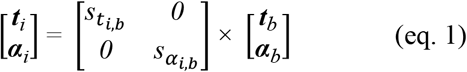

where ***t***_*i*_ and ***t***_*b*_ represent the time vectors, ***a***_*i*_ and ***a***_*b*_ the vectors with steering wheel angles, and *s_t_* and *s_a_* the scale factors for the time vector and the vector with steering wheel angles, respectively. To fit the scale factors, the data from the baseline trial and the trial of interest were first resampled to have matching lengths (i.e., the trial with the least samples was resampled using linear interpolation to have as many samples as the longer trial). Subsequently, scale factors were fitted by minimizing the combined sum of squared errors using the fminsearch function in MATLAB. For each trial, the fitted scale factors relative to the three baseline trials with a corresponding target distance and direction were averaged.

We additionally assessed skewness of the time traces of the steering wheel angle as a function of time by calculating Bowley’s coefficient of skewness for each trial *i*:

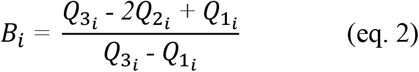

where *B* represents the skewness coefficient, and *Q_1_, Q_2_,* and *Q_3_* represent the times at which 25%, 50%, and 75% of the total distance travelled during the trial was covered, respectively. The skewness coefficients were baseline corrected by subtracting the average of the baseline trials with a corresponding target distance and direction. Negative skewness coefficients represent left-skewed steering profiles relative to baseline; positive skewness coefficients represent right-skewed steering profiles relative to baseline.

#### Statistics

Statistical analyses were done in R (version 4.0.1; see Ref. 28) using the package ez (version 4.4-0; see Ref. 29). Results were considered significant if the p-value was smaller than 0.05. To characterize baseline performance, we examined the average displacement error, movement duration and the maximum absolute steering wheel angle across the baseline trials (trials 73-90) with a mixed factorial ANOVA with condition (Motion and Stationary) as between-subject factor and target distance (20, 30 and 40 cm) and target direction (leftward and rightward) as within-subject factors. The results were adjusted according to the Greenhouse-Geisser correction in case of violations of sphericity. We report the generalized eta squared 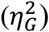 as a measure of the effect size (30).

To assess differences in changes in steering behavior in response to the two gain changes, we compared the behavior on trial 90 and trial 91 (high-to-low gain change) and on trial 162 and trial 163 (low-to-high gain change). We examined the relative displacement error, the two scale factors and the skewness coefficient using a mixed factorial ANOVA with condition (Motion and Stationary) as between-subject factor and gain change (high-to-low and low-to-high) as within-subject factor. Data from one participant in the Stationary condition were excluded from the analyses due to a trial rejection around the low-to-high gain change.

## Results

We created a motor signal of cognitive nature, enacted through a steering movement, and contrasted the learning of the relationship between the steering movement and a whole-body translation (Motion condition) with the translation of an object outside the body (Stationary condition). We focused on the availability of online vestibular feedback, allowing participants in the Motion condition to make online adjustments to their steering behavior in response to changes in the control dynamics.

Figure 2A shows the mean displacement error across participants as a function of the trial block per target distance and condition, pooled across target directions. Displacement errors across the baseline trials (trial blocks 13-15) were similar across conditions (*F*_1,28_ = 1.83, *p* = .187, 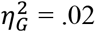) and target directions (*F*_1,28_ = 1.97, *p* = .172, 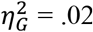), but varied across target distances (*F*_1.62,45.41_ = 33.95, *p* < .001, 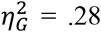). Participants were most accurate on baseline trials with an intermediate target distance (mean ± SD; 30 cm: 0.60 ± 3.94 cm), overshot the target location on trials with a small target distance (20 cm: 2.81 ± 3.07 cm), and undershot the target location on trials with a large target distance (40 cm: −3.18 ± 5.34 cm). Displacement errors across the baseline trials thus showed a range effect (31, 32). In the Stationary condition, participants undershot and overshot the target location shortly after the high-to-low and low-to-high gain change, respectively, irrespective of the target distance. However, the gain changes did not seem to influence the displacement error in the Motion condition. As this apparent lack of an effect of the gain changes in the Motion condition might be due to the low temporal resolution (trials were averaged across all six trials composing a trial block), we will refrain from statistics here. We will zoom in on the effect of the gain changes on the level of single trials later on.

**Figure 2.**
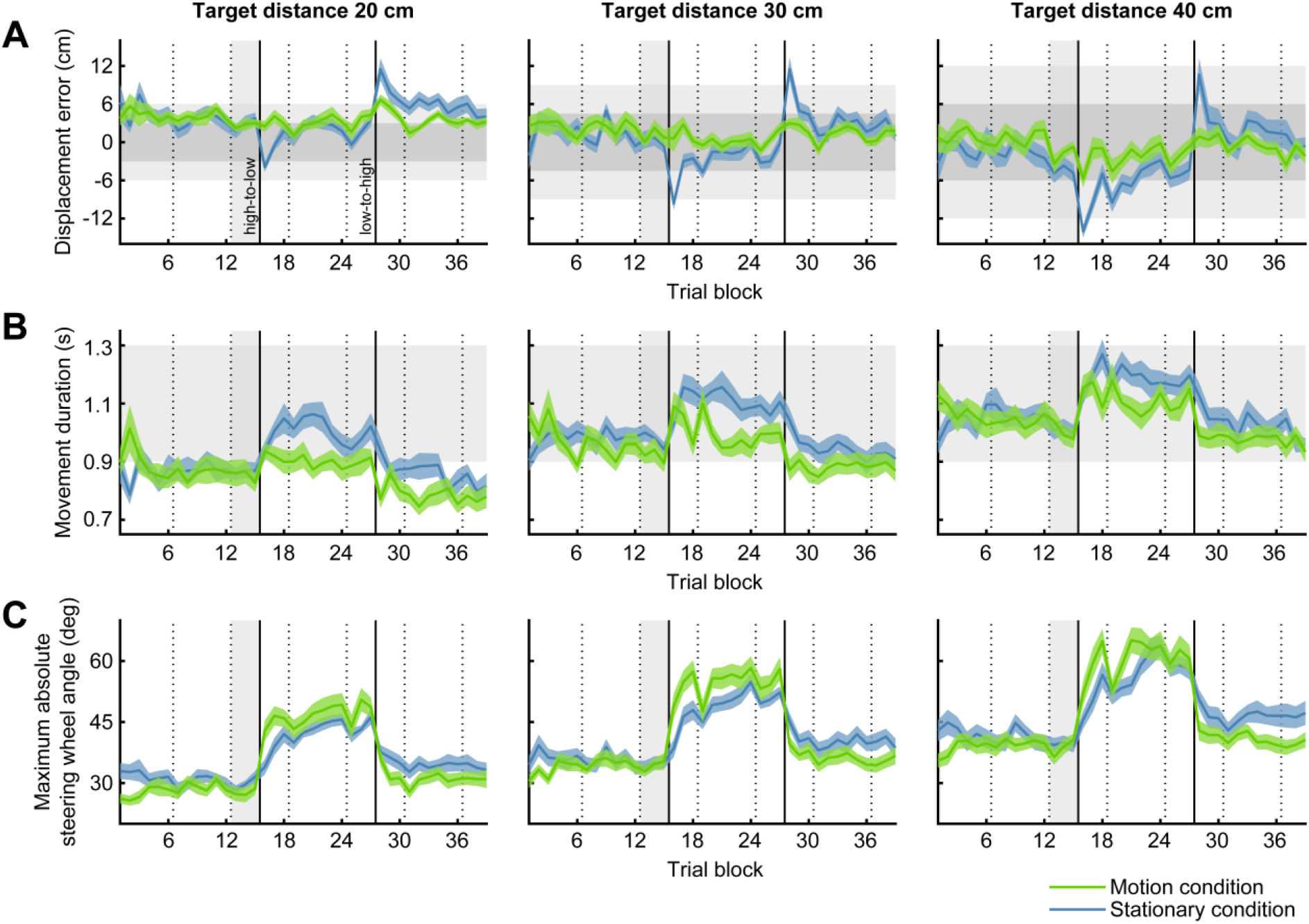
Displacement error, movement duration and maximum absolute steering wheel angle. A) Mean displacement error across participants as a function of trial block grouped based on target distance (panels) and experimental condition (colored lines). Displacement errors have been averaged across leftward and rightward displacements within a trial block. Negative numbers represent undershoots; positive numbers represent overshoots. Colored shaded areas represent between-subjects SEM. Horizontal dark and light gray bands show the range of displacement errors for which participants received 2 points and 1 point, respectively. Dashed vertical lines represent breaks, and solid vertical lines represent changes in the gain between the steering wheel angle and the velocity of the sled (Motion condition) or the line cursor (Stationary condition). Vertical light gray bands show the baseline trial blocks (trial blocks 13-15). A mixed factorial ANOVA revealed a significant main effect of target distance on the baseline displacement error (*p* < .001; Motion condition: n = 15; Stationary condition: n = 15). B) Same configuration as in A, but with the mean movement duration across participants. Horizontal light gray bands show the 900 to 1300 ms window within which participants were encouraged to finish their movement. A mixed factorial ANOVA revealed a significant main effect of target distance on the baseline movement duration (*p* < .001). C) Same configuration as in A, but with the mean maximum absolute steering wheel angle across participants. A mixed factorial ANOVA revealed a significant main effect of target distance on the baseline maximum absolute steering wheel angle (*p* < .001), as well as a significant interaction effect between target direction and experimental condition *(p* = .026, not visible in the figure).

The apparent lack of an effect of the gain changes on the displacement error in the Motion condition could also suggest that participants used the online vestibular feedback to make within-trial adjustments to their steering movement. These within-trial adjustments are likely to be reflected in the duration of the movement and the angle of the steering wheel. The velocity of the sled is directly affected by the gain changes, causing a sensory prediction error if an internal model has been developed, and both the movement duration and the steering wheel angle can be adjusted in response to this error. Figure 2B shows the mean movement duration across participants as a function of the trial block per target distance and condition, pooled across target directions. Movement duration across the baseline trials was similar across conditions (*F*_1,28_ = 0.58, *p* = .454, 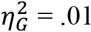) and target directions (*F*_1,28_ = 0.96, *p* = .337, 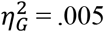), but varied across target distances (*F*_1.53,42.71_ = 46.57, *p* < .001, 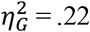). Participants took more time for the movement the longer the target distance (20 cm: 859 ± 125 ms; 30 cm: 947 ± 112 ms; 40 cm: 1006 ± 111 ms). Overall, the baseline movement duration was at the lower end of the imposed window from 900 to 1300 ms (Motion condition: 925 ± 147 ms; Stationary condition: 949 ± 111 ms). In the trial block after the high-to-low gain change, the movement duration increased immediately in the Motion condition across all target distances, followed shortly by the Stationary condition. Movement duration remained elevated, with a larger overall increase for the Stationary condition. In the trial block after the low-to-high gain change, movement duration immediately returned to baseline values in the Motion condition, whereas movement duration decreased a little more gradually in the Stationary condition.

Figure 2C shows the mean maximum absolute steering wheel angle across participants as a function of the trial block per target distance and condition, pooled across target directions. The maximum absolute steering wheel angle across the baseline trials was similar across conditions (*F*_1,28_ = 0.43, *p* = .520, 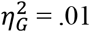) and target directions (*F*_1,28_ = 1.57, *p* = .221, 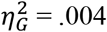), but varied across target distances (*F*_1.74,48.72_ = 117.34, *p* < .001, 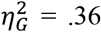). Participants increased the maximum angle with increasing target distances (20 cm: 28.74 ± 5.23 deg; 30 cm: 34.49 ± 5.60 deg; 40 cm: 39.09 ± 6.55 deg). We additionally found a small but significant interaction effect between target direction and condition (*F*_1,28_ = 5.55, *p* = .026, 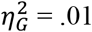). This interaction effect seems to be driven by a higher mean maximum absolute steering wheel angle for leftward than rightward displacements across the baseline trials in the Stationary condition (leftward: 35.75 ± 6.92 deg; rightward: 33.62 ± 6.54 deg), whereas the angle was similar across directions in the Motion condition (leftward: 33.19 ± 7.20 deg; rightward: 33.84 ± 7.95 deg). In the trial block after the high-to-low gain change, the maximum absolute steering wheel angle increased in the Motion condition across all target distances. The maximum absolute steering wheel angle remained relatively high until the low-to-high gain change, after which it decreased rapidly. In the Stationary condition, the maximum absolute steering wheel angle also increased and decreased after the high-to-low and low-to-high gain change, respectively, but more gradually.

To be able to inspect the effect of the gain changes at a high temporal resolution of single trials, while taking the semi-random trial order into account, we computed the relative displacement error as the ratio of the displacement error and the target distance. Figure 3A shows the relative displacement error across all trials, separately for the Motion condition and the Stationary condition. While the relative displacement error straddled closely around zero in the Motion condition, also after the gain changes, this was not the case in the Stationary condition, where there are clear deviations following the gain changes. Figure 3B illustrates the difference in the relative displacement error between the first trial after and the last trial before the gain changes, showing a significant main effect of the gain change (*F*_1,27_ = 57.54, *p* < .001, 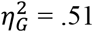) and a significant interaction effect between the gain change and the condition (*F*_1,27_ = 23.89, *p* < .001, 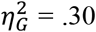). This indicates that the gain changes affected the relative displacement error differently across conditions, with larger relative errors in the Stationary condition (high-to-low: −0.53 ± 0.28; low-to-high: 0.85 ± 0.52) than in the Motion condition (high-to-low: −0.11 ± 0.32; low-to-high: 0.18 ± 0.50).

**Figure 3.**
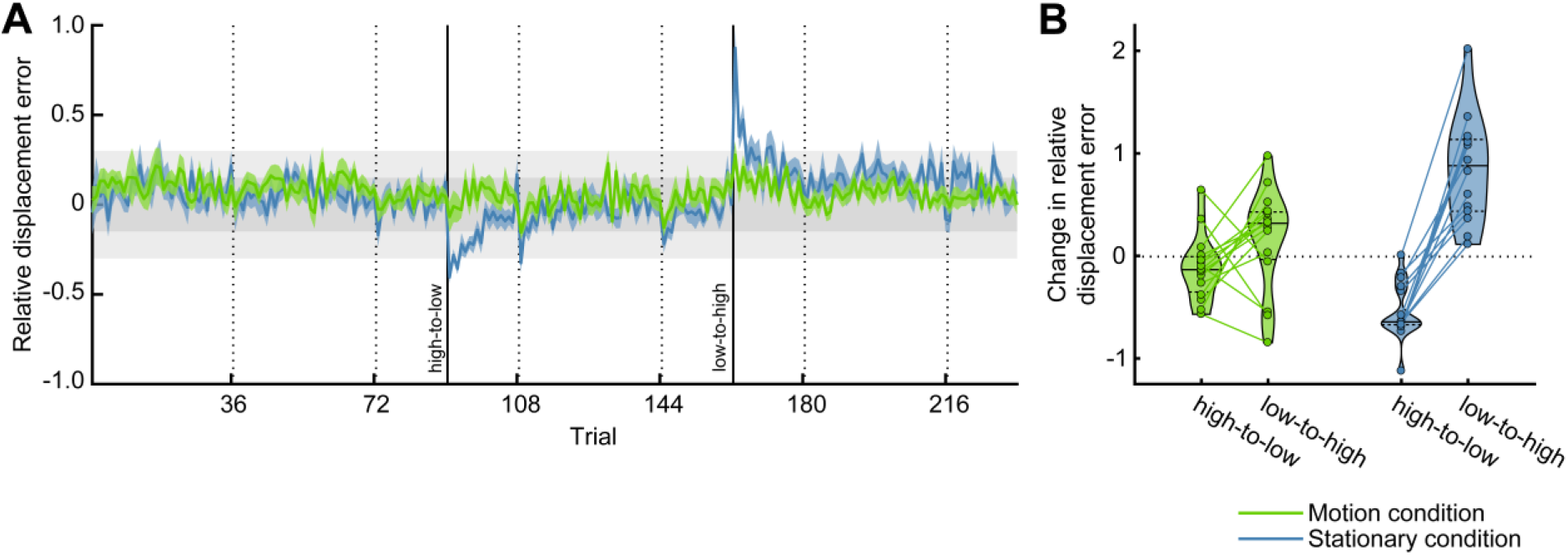
Relative displacement error. A) Mean relative displacement error across participants as a function of trial grouped based on experimental condition (colored lines). Relative displacement error was computed as the ratio of the displacement error and the target distance. Colored shaded areas represent between-subjects SEM. Dark and light gray bands show the range of displacement errors for which participants received 2 points and 1 point, respectively. Dashed vertical lines represent breaks, and solid vertical lines represent changes in the gain between the steering wheel angle and the velocity of the sled (Motion condition) or the line cursor (Stationary condition). B) Mean difference in relative displacement error between the first trial after and the last trial before the gain changes (high-to-low: trial 91 - trial 90; low-to-high: trial 163 - trial 162) across participants. Violin shape outlines show the kernel density estimates of the individual participant data points (colored dots connected by colored lines). Solid and dashed horizontal lines within the violin shapes represent the median and interquartile range, respectively. A mixed factorial ANOVA revealed a significant interaction effect between the gain change and the experimental condition (*p* < .001; Motion condition: n = 15; Stationary condition: n = 14), as well as a significant main effect of the gain change (*p* < .001).

The observation that the relative displacement error was virtually constant across gain changes in the Motion condition, also on the level of single trials, suggests that participants indeed used the online vestibular feedback to make within-trial adjustments to their steering movement, as described above. Figure 4 illustrates these within-trial adjustments in response to the gain changes. Within the first trial after the high-to-low gain change (trial 91, solid light green line), participants in the Motion condition increased the duration and the absolute steering wheel angle of their steering movement relative to the previous baseline trial (trial 90, dashed light green line) to compensate for the lower gain. Participants increased the absolute steering wheel angle further in later trials with this gain, as shown in the last trial with the low gain (trial 162, dashed medium green line). Conversely, within the first trial after the low-to-high gain change (trial 163, solid medium green line), participants decreased the duration and the absolute steering wheel angle again to compensate for the higher gain. At the end of the experiment (trial 234, dashed dark green line), participants returned to their baseline steering behavior. In the Stationary condition, no online feedback was available, and participants could thus not have been aware of the gain changes during the trials immediately after. This is also reflected in their behavior: the duration and the absolute steering wheel angle of their steering movement remained similar across the high-to-low gain change (trial 90 and 91, light blue lines) and the low-to-high gain change (trial 162 and 163, medium blue lines). However, participants did adjust their steering behavior in later trials with this gain, as shown by an increase in the duration and the absolute steering wheel angle from the first to the last trial with the low gain (trial 91 and 162, solid light blue line and dashed medium blue line, respectively) and a decrease in the duration and the absolute steering wheel angle back to baseline values from the first trial with the high gain to the last trial of the experiment (trial 163 and 234, solid medium blue line and dashed dark blue line, respectively).

**Figure 4.**
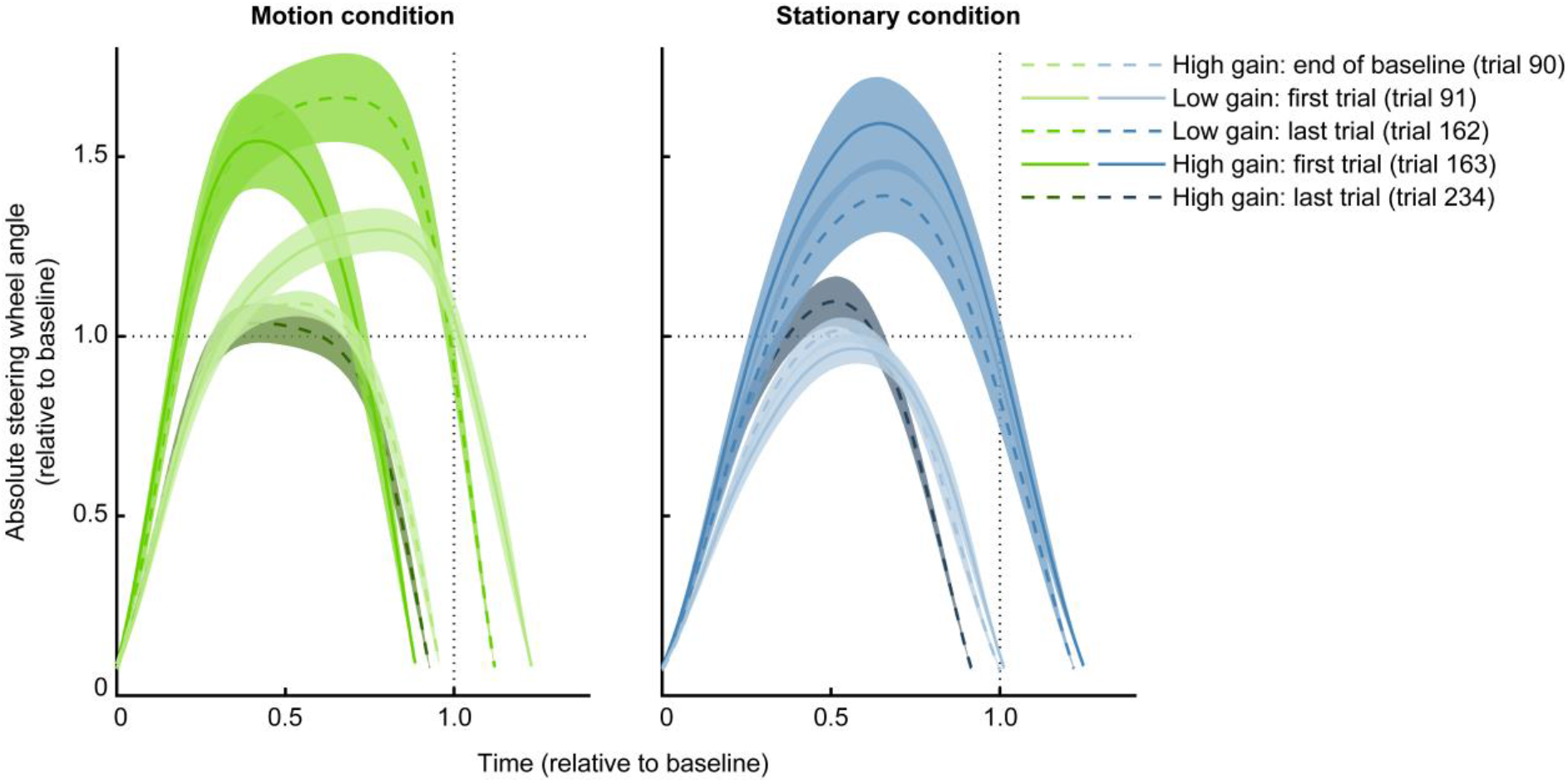
Steering behavior around the gain changes. Average absolute steering wheel angle as a function of time across participants grouped based on experimental condition (panels) for five key trials in the experiment, i.e., before and after the changes in gain between the steering wheel angle and the velocity of the sled (Motion condition) or the line cursor (Stationary condition). Values were normalized relative to baseline; time values equal to zero represent movement onset, time values equal to one represent the mean movement duration across baseline trials (dashed vertical lines), and absolute steering wheel angles equal to one represent the mean maximum absolute steering wheel angle across baseline trials (dashed horizontal lines). Lines with the same color represent the last trial before (dashed line) and the first trial after (solid line) one of the two gain changes (high-to-low: light colors; low-to-high: medium colors). Dashed dark colored lines represent the last trial of the experiment. Note that trial 90, represented by the dashed light colored lines, belongs to the baseline. Colored shaded areas represent between-subjects SEM.

We quantified the within-trial adjustments in response to the gain changes in both conditions at a high temporal resolution of single trials by scaling of the time-axis and steering wheel angle-axis relative to the baseline (trials 73-90). With this linear transformation, the axes of the trial of interest are independently stretched and compressed to match the baseline trial. Figures 5A and 5B show the fitted scale factors for the time-axis, describing the movement duration, and the steering wheel angle-axis, respectively, across all trials. In line with Figure 4, both the movement duration and steering wheel angle increased relative to baseline immediately after the high-to-low gain change and decreased immediately after the low-to-high gain change in the Motion condition. Similar patterns were observed for the Stationary condition, albeit with a one-trial delay and slower changes in behavior. Participants in both conditions continued to adjust their steering behavior in response to the gain changes across trials. This is most clearly visible after the high-to-low gain change: after the immediate increase in movement duration and steering wheel angle, participants continued to increase the steering wheel angle across trials while decreasing the movement duration. The latter is not surprising, as we encouraged participants to finish their steering movement within a time window from 900 to 1300 ms, and thereby indirectly encouraged them to adjust the steering wheel angle instead of the movement duration.

**Figure 5.**
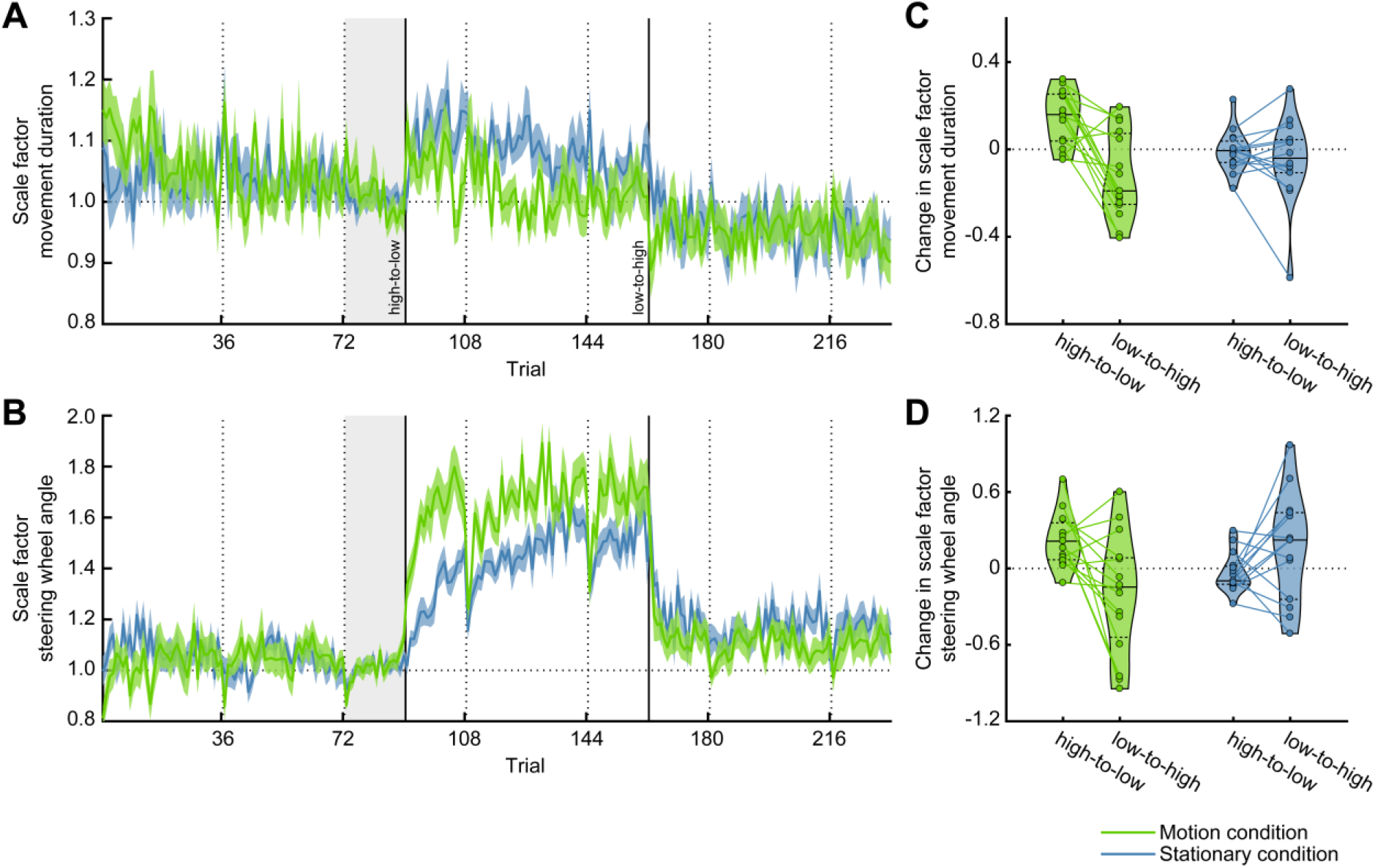
Scale factors movement duration and steering wheel angle. A) Mean movement duration scale factor across participants as a function of trial grouped based on experimental condition (colored lines). Scale factors were fitted relative to the baseline trials (trials 73-90, vertical light gray band) with a corresponding target distance and direction. Colored shaded areas represent between-subjects SEM. Dashed vertical lines represent breaks, and solid vertical lines represent changes in the gain between the steering wheel angle and the velocity of the sled (Motion condition) or the line cursor (Stationary condition). B) Same as in A, but with the steering wheel angle scale factor. C) Mean difference in the movement duration scale factor between the first trial after and the last trial before the gain changes (high-to-low: trial 91 - trial 90; low-to-high: trial 163 - trial 162) across participants. Violin shape outlines show the kernel density estimates of the individual participant data points (colored dots connected by colored lines). Solid and dashed horizontal lines within the violin shapes represent the median and interquartile range, respectively. A mixed factorial ANOVA revealed a significant interaction effect between the gain change and the experimental condition (*p* = .003; Motion condition: n = 15; Stationary condition: n = 14), as well as a significant main effect of the gain change (*p* < .001). D) Same as in C, but with the steering wheel angle scale factor. A mixed factorial ANOVA revealed a significant interaction effect between the gain change and the experimental condition (*p* = .003).

Figure 5C illustrates the difference in the movement duration scale factor between the first trial after and last trial before the gain changes, showing a significant main effect of the gain change (*F*_1,27_ = 14.56, *p* < .001, 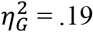) and a significant interaction effect between the gain change and the condition (*F*_1,27_ = 10.34, *p* = .003, 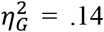). This indicates that the gain changes affected the scale factor differently across conditions, with larger changes in the movement duration in the Motion condition (high-to-low: 0.15 ± 0.12; low-to-high: −0.12 ± 0.20) than in the Stationary condition (high-to-low: −0.01 ± 0.09; low-to-high: −0.05 ± 0.20). Figure 5D illustrates the difference in the steering wheel angle scale factor across the trials just before and after the gain changes, showing a significant interaction effect between the gain change and the condition (*F*_1,27_ = 10.66, *p* = .003, 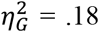). Similar to the effects described for the movement duration above, the changes in the steering wheel angle were larger in the Motion condition (high-to-low: 0.22 ± 0.21; low-to-high: −0.20 ± 0.47) than in the Stationary condition (high-to-low: −0.04 ± 0.15; low-to-high: 0.17 ± 0.42). Here, no significant main effect of the gain change was observed (*F*_1,27_ = 1.39, *p* = .249, 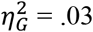).

These results show that participants in the Motion condition used the online vestibular feedback to change their steering behavior within the first trial after the gain changes. We next examined the skewness of the observed steering profiles. If participants in the Motion condition developed an internal model of the mapping between the steering movement and the vestibular reafference and a sensory prediction error is detected early in the motion after a gain change, a rapid correction is expected, causing a skew of the steering profile. We computed Bowley’s skewness coefficient for each trial. This skewness coefficient provides information about how the distance travelled is distributed across the trial duration. Figure 6A shows the mean skewness coefficient across participants as a function of trial number, separately for the Motion and the Stationary condition. In the Motion condition, the skewness coefficient decreased after the high-to-low gain change, indicating a left-skewed steering profile (i.e., the increase of the absolute steering wheel angle was slower than the decrease, see also trial 91 in Figure 4). After the low-to-high gain change, the skewness coefficient increased, indicating a right-skewed steering profile (i.e., the increase of the absolute steering wheel angle was faster than the decrease, see also trial 163 in Figure 4). Both changes in the skewness coefficient were short-lasting and did not persist across the trials following the first trial after the gain changes. In the Stationary condition, skewness coefficients remained rather constant across trials and gain changes. Figure 6B illustrates the difference in the skewness coefficient across the trials just before and after the gain changes, showing a small but significant interaction effect between the gain change and the condition (*F*_1,27_ = 5.54, *p* = .026, 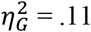). This indicates that the gain changes affected the skewness coefficient differently across conditions, with more skewed steering profiles in the Motion condition (high-to-low: −0.017 ± 0.033; low-to-high: 0.020 ± 0.034) than in the Stationary condition (high-to-low: −0.001 ± 0.015; low-to-high: −0.005 ± 0.033). No significant main effect of the gain change was found (*F*_1,27_ = 3.83, *p* = .061, 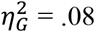).

**Figure 6.**
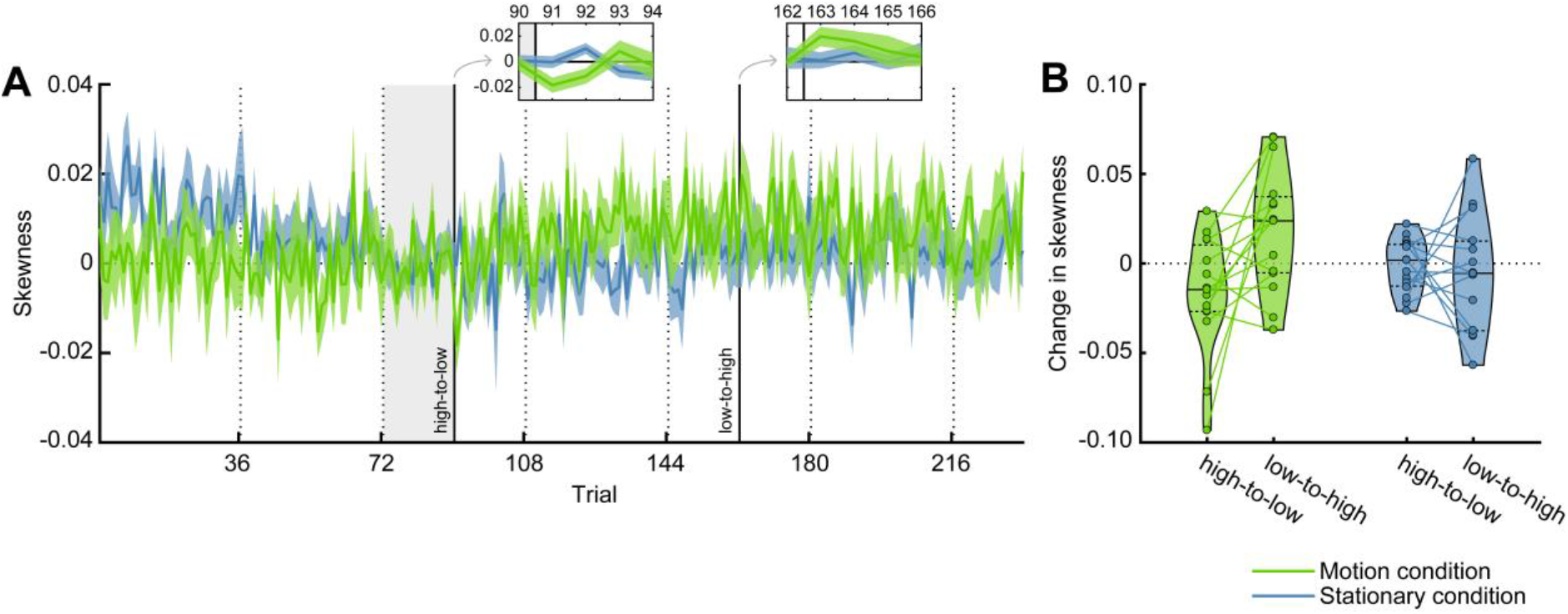
Skewness. A) Mean skewness coefficient across participants as a function of trial grouped based on experimental condition (colored lines). Skewness coefficients were baseline corrected by subtracting the average across the baseline trials (trials 73-90, vertical light gray bands) with a corresponding target distance and direction. Negative skewness coefficients represent left-skewed steering profiles; positive skewness coefficients represent right-skewed steering profiles. Colored shaded areas represent between-subjects SEM. Dashed vertical lines represent breaks, and solid vertical lines represent changes in the gain between the steering wheel angle and the velocity of the sled (Motion condition) or the line cursor (Stationary condition). Insets show the zoomed views of the trials before and after the gain changes. B) Mean difference in the skewness coefficient between the first trial after and the last trial before the gain changes (high-to-low: trial 91 - trial 90; low-to-high: trial 163 - trial 162) across participants. Violin shape outlines show the kernel density estimates of the individual participant data points (colored dots connected by colored lines). Solid and dashed horizontal lines within the violin shapes represent the median and interquartile range, respectively. A mixed factorial ANOVA revealed a significant interaction effect between the gain change and the experimental condition (*p* = .026; Motion condition: n = 15; Stationary condition: n = 14).

## Discussion

Participants were tested in a naturalistic self-motion task in which they actively controlled their own body motion on a motion sled, while traversing to a remembered target location in darkness (Motion condition). The goal was to examine whether participants estimated their self-motion based on the vestibular signals resulting from their body motion and anticipated the vestibular reafference using an internal forward model with the steering efference copy as input. To test whether participants predicted the vestibular reafference and integrated their predictions with the actual vestibular feedback, we unexpectedly changed the gain between the steering movement and the sled motion twice during the experiment and recorded participants’ changes in steering behavior. We compared their steering behavior with that of participants who controlled a line cursor instead of their own body motion, and thus did not have access to online vestibular feedback (Stationary condition).

In the Motion condition, the sudden gain changes did not result in systematic changes in displacement errors (Fig. 2 and 3). Instead, we observed within-trial changes in steering behavior immediately after the gain changes; participants increased and decreased the movement duration and the steering wheel angle to compensate for the high-to-low and the low-to-high gain changes, respectively (Fig. 4 and 5). These within-trial adjustments, resulting in skewed steering profiles (Fig. 6), show that participants continuously monitored the available vestibular feedback to keep track of their self-motion and compared it to the expected vestibular reafference. Additionally, participants continued to revise the movement duration and steering wheel angle in subsequent trials with the new gain, gradually improving their adaptation to the new control dynamics (i.e., revise the movement duration and the steering wheel angle to be able to adhere to the imposed movement duration). This shift from fast and reactive changes in behavior to more tactful and planned changes suggests that participants built and updated an internal model of the steering signal and the associated self-motion based on the online vestibular feedback.

In contrast, in the Stationary condition, the gain changes resulted in substantially increased displacement errors (Fig. 2 and 3). This is not surprising; participants assigned to this condition found out about the gain changes at the earliest at the end of the first trial after the gain changes, based on the visual feedback about their displacement error. Across trials, however, these participants adjusted their steering behavior based on this feedback to compensate for the gain changes; after the gain changes they gradually changed the movement duration and the steering wheel angle, without changing the skewness of their steering movement (Fig. 4, 5 and 6), causing the displacement error to decrease. Overall, the results suggest that also the participants in the Stationary condition built an internal model, illustrated by their overall ability to perform the task (i.e., the displacement error at the end of the baseline did not differ from the Motion condition), and employed a feedforward control strategy to gradually improve their performance across trials after the gain changes based on the visual feedback at the end of the trial.

Could participants in the Motion condition have performed the task without forming an internal model (and thus without online predictions of the self-motion, i.e., the vestibular reafference)? By integrating the vestibular information relating to the velocity of the sled over time – as in models of path integration (33) – participants could have kept track of the position of the sled in space, and stopped the sled when the required travel distance, specified by the target, was reached. However, the fast changes in steering behavior in response to the gain changes, as also shown by the changes in the scale factors and skewness coefficient describing the steering profiles (Fig. 4, 5 and 6), suggest that participants had some expectations relating to the velocity of the sled. So, the tentative explanation of our results is that participants are able to generate predictions about the vestibular feedback based on artificial motor signals (i.e., the steering movement) and compare these predictions to the actual online vestibular feedback in order to estimate their self-motion (16, 17). These computations are similar to those underlying the perception of true active self-motion, and our results therefore suggest that artificial signals, such as the steering motor signal, can serve as an efference copy that can be integrated in self-motion perception.

The present study builds on experiments in both self-motion perception and motor learning. Motor learning experiments, such as force field experiments during which reaching movements are perturbed by forces applied to the arm, have shown that participants adapt to new but sustained environments faster if online (visual) feedback is available (34, 35). Additionally, even when the environment is completely unpredictable and changes from trial to trial, participants have been shown to be able to use online feedback to adapt by adjusting their behavior (36). Our results are in line with these observations; after the unexpected gain changes, participants in the Motion condition, who had online vestibular feedback, adjusted their steering behavior faster than participants in the Stationary condition. Of note, reaching movements are often ballistic, with movement durations around 600 ms, and are therefore likely to depend to a large extent on feedforward processes. The steering movements in the current experiment were slower, with movement durations around 900 ms, and there might therefore have been even more time for online adjustments.

Our study is one among the few recent studies that tested self-motion perception under a direct coupling between the actions of the participants and the sensory feedback (25–27). These recent experiments imposed fewer artificial constraints than the traditional open-loop psychophysical paradigms on self-motion perception (e.g., 37–41). In the previous psychophysical experiments, the perception of self-motion was assessed by responses on a two-alternative forced choice task, allowing to estimate how the brain weighs and adapts to sensory cues during the motion in the absence of changing motor cues (41, 42). While these experiments have led to important advances in the self-motion perception field, they cannot inform us how cues with time-varying noise levels are integrated over longer periods of time or which utility functions and task dependencies guide the naturalistic closed-loop navigation behavior.

In a recent study by Stavropoulos et al. (27), human participants controlled their self-displacement on a motion platform to navigate to a target with and without the presence of concurrent optic flow. They found that steering behavior in darkness was biased (i.e., participants undershot the target location), and therefore concluded that participants could not accurately estimate their self-motion and update their internal model based on the vestibular cues alone. This conclusion differs from the present results, but could be explained by differences in the experimental design. More specifically, their participants did not receive any performance-related feedback, and the control dynamics of the motion platform changed from trial to trial, both of which may have kept participants from building an accurate internal model of the mapping between the steering movement and the vestibular feedback.

The rapid changes in steering behavior in response to the gain changes suggest that our participants had some expectations about the ensuing self-motion. Further support for this notion comes from previous studies showing that, during passive but predictable self-motion, the effects of the self-motion are anticipated. For example, during passively-induced angular (see Ref. 43 for a review) and linear whole-body displacements (44), participants were able to anticipate and counteract the inertial forces exerted on the arm, resulting in accurate goal-directed reaching movements. Also Prsa et al. (18) showed that passive angular displacement estimates in human participants were biased towards the average over a block of random displacement magnitudes, suggesting that participants built up some expectations about the vestibular input.

Roy and Cullen (21) have shown that neurons in the vestibular nuclei (VN) of monkeys respond similarly during steering-controlled and passively-induced self-motion. Under the assumption that the firing rates of neurons in the VN reflect sensory predictions errors (16, 45), this suggests that no steering-related predictions about the vestibular reafference are made in the VN. Even though the vestibular cerebellum is often suggested to house the internal model for self-motion estimation because of its projections to the vestibular nuclei (46), the internal model of the mapping between steering movements and self-motion seems located on a more downstream level within the vestibular processing pathway (25). This is in line with observations during the processing of the visual reafference of steering movements; Page and Duffy (22) reported that neurons in the medial superior temporal area in monkeys responded differently to optic flow cues resulting from steering movements compared to passive viewing of the same optic flow cues.

The use of artificial signals in self-motion perception is currently exploited in the development of vestibular implants for patients with a vestibular deficit (47, 48). These vestibular implants electrically stimulate the vestibular nerve in a biomimetic way and provide patients with artificial vestibular feedback. Similarly, these patients have been shown to benefit from tactile and auditory cues that provide information about the vestibular input through an arbitrary mapping (see Ref. 47 for a review). This mapping has to be learned, similar to the gain in the present study, and the learning of such a mapping has even been extended to augmenting perception in healthy human subjects by adding an extra “vestibular” sense (i.e., head orientation relative to the geomagnetic North) (49). Altogether, these experiments show that participants can learn the mapping between an artificial sensory feedback signal and their self-motion, similar to the artificial motor signal used in the current study.

## Author contributions

M.J.L.H., L.P.J.S., R.J.B. and W.P.M. conceived and designed research; M.J.L.H. performed experiments; M.J.L.H., L.P.J.S., R.J.B. and W.P.M. analyzed data; M.J.L.H., L.P.J.S., R.J.B. and W.P.M. interpreted results of experiments; M.J.L.H. prepared figures; M.J.L.H., L.P.J.S., R.J.B. and W.P.M. drafted manuscript; M.J.L.H., L.P.J.S., R.J.B. and W.P.M. edited and revised manuscript; M.J.L.H., L.P.J.S., R.J.B. and W.P.M. approved final version of manuscript.

## Data availability statement

Upon publication, all data and code will be made publicly available via the persistent identifier currently reserved for this collection: https://doi.org/10.34973/74t0-w530.

